# Intraspecific *de novo* gene birth revealed by presence–absence variant genes in *Caenorhabditis elegans*

**DOI:** 10.1101/2021.09.24.461648

**Authors:** Bo Yun Lee, Jun Kim, Junho Lee

## Abstract

Genes embed their evolutionary history in the form of various alleles. Presence–absence variants (PAVs) are extreme cases of such alleles, where a gene present in one haplotype does not exist in another. Since PAVs may result from either birth or death of a gene, PAV genes and their alternative alleles, if available, can represent a basis for rapid intraspecific gene evolution. Here, we traced a possible evolution of PAV genes in the PD1074 and CB4856 *C. elegans* strains as well as their alternative alleles found in other 14 wild strains, using long-read sequencing technologies. We updated the CB4856 genome by filling 18 gaps and identified 50 novel genes and 7,460 novel isoforms from both strains. We verified 328 PAV genes, out of which 48 were *C. elegans*-specific. Among these possible newly-born genes, 13 had alternative alleles in other wild strains and, in particular, alternative alleles of three genes showed signatures active transposons. Alternative alleles of four other genes showed another type of signature reflected in accumulation of small insertions or deletions. Our results exemplify that research on gene evolution using both species-specific PAV genes and their alternative alleles is expected to provide new perspectives for how genes evolve.

## INTRODUCTION

Genetic variation shapes sequences of genes from their birth to their death. Gene duplication and divergence are the most representative examples by which one functional gene is duplicated in multiple copies through segmental or whole-genome duplication, so that copied genes acquire divergent sequences and new functions (1–4). A new gene can also be born in a non-genic region through a process in which non-genic transcripts acquire open reading frames and functions, which is called *de novo* gene birth (4–8). Conversely, genes may disappear as genetic variation disrupts their functions or deletes the entire gene sequences, resulting in gene death (4). These concepts have advanced our understanding of how genes are newly born or vanished, but details of these events remain elusive in a short time scale, at the intraspecific level.

A type of genetic variants, presence–absence variants (PAVs), is represented by genes that are present in some genomes but absent in others within the same species. PAV genes might be fast-evolving genes, as they are able to reflect birth or death of genes (9). In addition, PAV genes may contribute to adaptation to changing environments, including pathogen infection, antitumor-agent synthesis or disease resistance in plants as well as immunity in animals (10–14). Recently, another case of gene evolution was also analysed using PAVs in several molluscs (15). Thus, PAVs may provide lines of evidence of how genes evolve, but some of their characteristics make it difficult to identify PAVs precisely. PAVs typically do not contain conserved or essential genes, as they should not have a disruptive function, thus homology-based gene prediction may fail to detect such orphan genes. High-quality genome assembly and evidence-based gene annotation methods can be employed, but stereotypical sequencing techniques are based on short-read sequencing technologies that sometimes result in fragmented genomes and do not cover full-length transcripts. Advances in long-read sequencing technologies resolve these limitations and make it possible to detect PAVs at the population level (16–18).

*Caenorhabditis elegans* is a model organism suitable to identify PAVs at the population level. It has a small genome size of about 100 Mb and hundreds of wild strains are isolated and cryopreserved (19,20). In a previous study, genes absent in 12 wild strains, including CB4856, but present in the reference strain were discovered; however, genes present specifically in these strains could not be identified at the sequence level, since the authors used a hybridisation-based method (21). Long-read sequencing-based high-quality genomic resources are resolving these technical limitations, thus providing the opportunity to better understand PAVs in *C. elegans* populations. For instance, a clonal selection-descendant strain of the N2 reference strain, PD1074, was established to eliminate mutations accumulated in N2 and its high-quality genome containing only two gaps was completed (22). Moreover, the genome of CB4856, one of genetically divergent strains from the reference strain, was completed at the pseudo-chromosome level and high-quality genomes of 14 wild strains were also assembled (23,24). It further clears the way to determine which genes represent PAV genes in *C. elegans* and to identify their alternative alleles, which could lead to clarifying the processes of possible gene birth or death.

Here, we analyse *C. elegans*-specific PAV genes and their alternative alleles in order to infer possible scenarios of gene evolution within a given species. We report the identification of novel genes and transcripts from both PD1074 and CB4856 genomes by using a long-read RNA sequencing technology. We then show that *C. elegans*-specific PAV genes and their alternative alleles identified from 14 *C. elegans* wild strains manifest snapshots of gene evolution such as complex small insertions and deletions (indels), active transposon signatures, and gene duplication and divergence signatures. We discuss that the analysis of alternative alleles of the PAV genes within a given species is a useful tool to understand clues of gene evolution that occurs in a short period of time.

## MATERIAL AND METHODS

### *C. elegans* strains and maintenance

*C. elegans* worms belonging to two strains, PD1074 and CB4856, were cultured under standard culture conditions.

### Genomic DNA extraction and ONT sequencing

Mixed-stage worms of the CB4856 strain grown at 20°C were harvested and washed 5 times with M9 buffer. Worms were lysed in Cell Lysis Solution from The Gentra Puregene^®^ Cell and Tissue Kit (Qiagen) with 0.1 mg/mL proteinase K and 1% β-mercaptoethanol at 55°C for 2 h. We purified DNA three times using phenol/chloroform extraction and ethanol precipitation coupled with phase-lock gel to minimise DNA shearing. DNA dissolved in TE buffer was treated with 10 μg/mL RNase for 2 h after the first extraction and precipitation step. DNA was treated with ONT SQK-LSK109 library preparation kit and the DNA library was sequenced using FLO-MIN106.

### Total RNA extraction and PacBio sequencing

Mixed-stage worms of both PD1074 and CB4856 strains were grown at 15°C, 20°C or 25°C. Each sample was separately harvested with M9 buffer. Worms were incubated in M9 with rotation for 30 minutes to remove bacteria in the gut and washed three times with M9. Worms about 200 μL were treated with 2 mL of Trizol solution and subsequently disrupted by six freeze-thaw cycles. RNA was extracted with chloroform/isopropanol precipitation. Each RNA sample was quantified and samples collected at three different temperatures of each strain were pooled in equal amounts. Macrogen conducted RNA library preparation and Iso-Seq using the PacBio Sequel System.

### Gap filling in the CB4856 genome

We conducted basecallling of long-read DNA sequencing reads and trimming of adapter sequences using Guppy basecaller (version 3.4.1) with the default setting. We extracted reads longer than 20 kb and aligned them to the CB4856 genome (23) using Minimap2 (version 2.17; *minimap2 -ax map-ont*) (25). The selected reads were then filtered based on their primary alignments using Samtools (version 1.9; *samtools view -f 0×10 -q 2* or *samtools view -q 2*) (26). The filtered reads were aligned to the CB4856 genome again using Minimap2 and gap-filling reads were confirmed by visualising their alignments using NUCmer and mummerplot from MUMmer package (version 4.0.0 beta2) (27) and Gnuplot (version 5.0, patch level 3). We collected the reads that aligned to both flanking contigs of gaps and manually replaced reference sequences with their corresponding gap-filling read sequences (**Supplementary Table S2**).

### Iso-Seq data processing

We used the IsoSeq package (version 3.3.0) to process Iso-Seq data (https://github.com/PacificBiosciences/IsoSeq_SA3nUP/). First we generated circular consensus sequences (CCS) using CCS (version 4.2.0; *ccs --min-rq 0.8 -min-passes 1)* and removed library-prep-primers, poly(A) tails and artificial concatemers using lima (version 1.11.0; *lima --isoseq --dumpclips --peek-guess*) and IsoSeq (version 3.3.0; *isoseq3 refine --require-polya)*. After clustering and polishing full-length reads using IsoSeq (version 3.3.0; *isoseq3 cluster --verbose --use-qvs*), we aligned high-quality full-length non-concatemer reads to the PD1074 genome (WS274) or the CB4856 genome using Minimap2 (version 2.17; *minimap2 -ax splice -uf --secondary=no -C5, -O6,24 -B4)* (25). Finally, we extracted 5’ non-degraded isoforms using cDNA_Cupcake programme (version 12.0.0, *collapse_isoforms_by_sam.py --dun-merge-5-shorter* and *filter_away_subset.py*; https://github.com/Magdoll/cDNA_Cupcake).

### Transferring gene annotations from the PD1074 genome to the CB4856 genome using LiftOver

To annotate genes of CB4856, we conducted a chain alignment process between the CB4856 genome and the PD1074 genome using the same-species liftover construction method (http://genomewiki.ucsc.edu/index.php/Same_species_lift_over_construction) from UCSC (28). Referring to the method of Yoshimura et al. (22), *targetChunkSize* and *queryChunkSize* were modified from 10,000,000 to 22,000,000 and a typographical error was corrected from ‘B=‘basename [ file]’ to ‘B=‘basename [ file]”. The PD1074 gene annotation contained genes that were lifted from the reference N2 gene annotation and genes that were predicted based on the PD1074 genome. We only transferred PD1074 gene annotations (WS274) that originated from the N2 genome as they were better validated (*liftOver -gff*).

### Annotation of isoforms obtained by Iso-Seq

We annotated 5’ non-degraded high-quality full-length isoforms using SQANTI3 (version 1.1; *sqanti3_qc.py --aligner_choice=minimap2 --fl_count*) (29). We used the PD1074 genome and its N2-origin gene annotations as well as the CB4856 genome and transferred gene annotations as references for the SQANTI3 analysis. Then, we categorised the annotated genes as follows: known genes with only known isoforms, known genes with novel isoforms, novel gene candidates and fusion or not curated genes.

### Validating novel genes and their protein identity

We built BLAST database of the PD1074 genome, the CB4856 genome, the N2 transcriptome (WS274; mRNA, ncRNA, pseudogenic and transposon transcripts), the PD1074 transcripts (WS274; we only used transcripts transferred from N2) and our PD1074 long-read transcripts. After identifying and masking out low complexity parts of the sequences using DustMasker (version 1.0.0; *dustmasker -infmt fasta -parse_seqids -outfmt maskinfo_asnl_bin*), we built BLAST database using Makeblastdb (version 2.7.1+; *makeblastdb -input_type fasta -dbtype nucl -parse_seqids*). We then filtered out novel gene candidates from the SQANTI3 results by searching their CCS read sequences in all databases stated above (BLASTn version 2.7.1+; default setting). Finally, the validated novel gene sequences were searched using the NCBI BLASTp database to determine their protein identity (https://blast.ncbi.nlm.nih.gov/Blast.cgi?PAGE=Proteins) and using Batch CD-search (https://www.ncbi.nlm.nih.gov/Structure/bwrpsb/bwrpsb.cgi) to identify any conserved domains (30).

### Identifying PAVs and determining their characteristics

Genomic sequences of the PD1074 strain annotated by SQANTI3, excluding transferred genes in the PD1074 gene annotations, were analysed in the CB4856 genome database using BLASTn (version 2.7.1 +; *blastn* for sequences longer than 50 bp or *blastn -task blastn-short* for sequences shorter than or equal to 50 bp) to identify PD1074-specific genes. CB4856-specific genes were identified by searching genomic sequences of the CB4856 SQANTI3 results in the PD1074 genome database. We considered as PAVs all genes from each strain that did not exhibit any significant identity in BLAST search results.

We searched these PAVs in the genomes of other 14 wild strains (DL238: GCA_016989505.1; ECA36: GCA_016989455.1, ECA396: GCA_016989385.1, JU310: GCA_016989275.1, JU1400: GCA_016989365.1, JU2526: GCA_016989285.1, EG4725: GCA_016989295.1, JU2600: GCA_016989245.1, MY2147: GCA_016989235.1, NIC2: GCA_016989145.1, NIC526: GCA_016989115.1, QX1794: GCA_016989095.1, MY2693: GCA_016989105.1 and XZ1516: GCA_016989125.1) (24). We first made each genome database for the 14 wild strains using Makeblastdb (version 2.7.1+; *makeblastdb -input_type fasta -dbtype nucl-parse_seqids*). After performing BLAST search (BLASTn version 2.7.1+; default setting), we classified our PAVs status in each strain according to their coverage in BLAST results for the corresponding strain as follows: no significant identity — absence, significant identity with coverage ≥ 90% — presence, in-between — alternative. Raw data are summarised in **Supplementary Table S11** and **S12**.

These strain-specific protein-coding genes were also analysed against the NCBi protein database and the Batch CD-search database to identify any similarity to known protein or domain sequences. We used only genes with e-value < 0.001 and categorised them as follows: genes that do not have significant identity with genes of any other species — *C. elegans*-specific genes, genes that have similarity only with genes of other *Caenorhabditis* species — *Caenorhabditis*-specific genes and genes that have similarity with genes of species in other animal phyla — conserved genes. For PD1074-specific genes, we also tested whether they have known RNAi phenotypes using the SimpleMine tool in WormBase (https://wormbase.org//tools/mine/simplemine.cgi) (31). The phenotypes were confirmed through literature search (32–38).

### Identifying alternative alleles in wild strains

Alternative alleles present in wild strains were identified by searching for presence alleles of our PAV genes in the genome assemblies of the 14 wild strains (24). The presence allele sequences included UTR regions. Because the searched sequences were partial in the 14 genome assemblies, we defined the alternative alleles as the searched sequences and their flanking sequences extended to the length of unsearched regions in the corresponding presence alleles. We obtained the coverage of each alternative allele by aligning their sequences to the N2 genome (WS279) using the BLAST/BLAT tool of WormBase (https://wormbase.org/tools/blast_blat) (39).

## RESULTS

### Updating the CB4856 genome with ONT long reads

Since precise identification of PAVs partly depends on the genome quality, we first updated the previously published CB4856 genome (23) by filling gaps between contigs with ONT long-read DNA sequencing data. We obtained 57-fold coverage of DNA sequences with a read-length N50 of 4,284 bp and a total read-length of 5.9 Gb (**Supplementary Table S1** and **Supplementary Figure S1A**). To maximise the benefits of long-read sequencing, we used only long reads with over 20-kb read-length and aligned the resulting 12,982 reads to the CB4856 genome (**Supplementary Figure S1B**). We found 27 reads that were able to connect two flanking contigs around the gaps (**Supplementary Table S1** and **S2** and **Supplementary Figure S2**) and filled 18 gaps with these reads. We updated the previous CB4856 genome composed of 76 contigs with a 103-Mb genome made of 54 contigs (**Table 1**). We used this updated genome for further analyses.

**Table 1.**
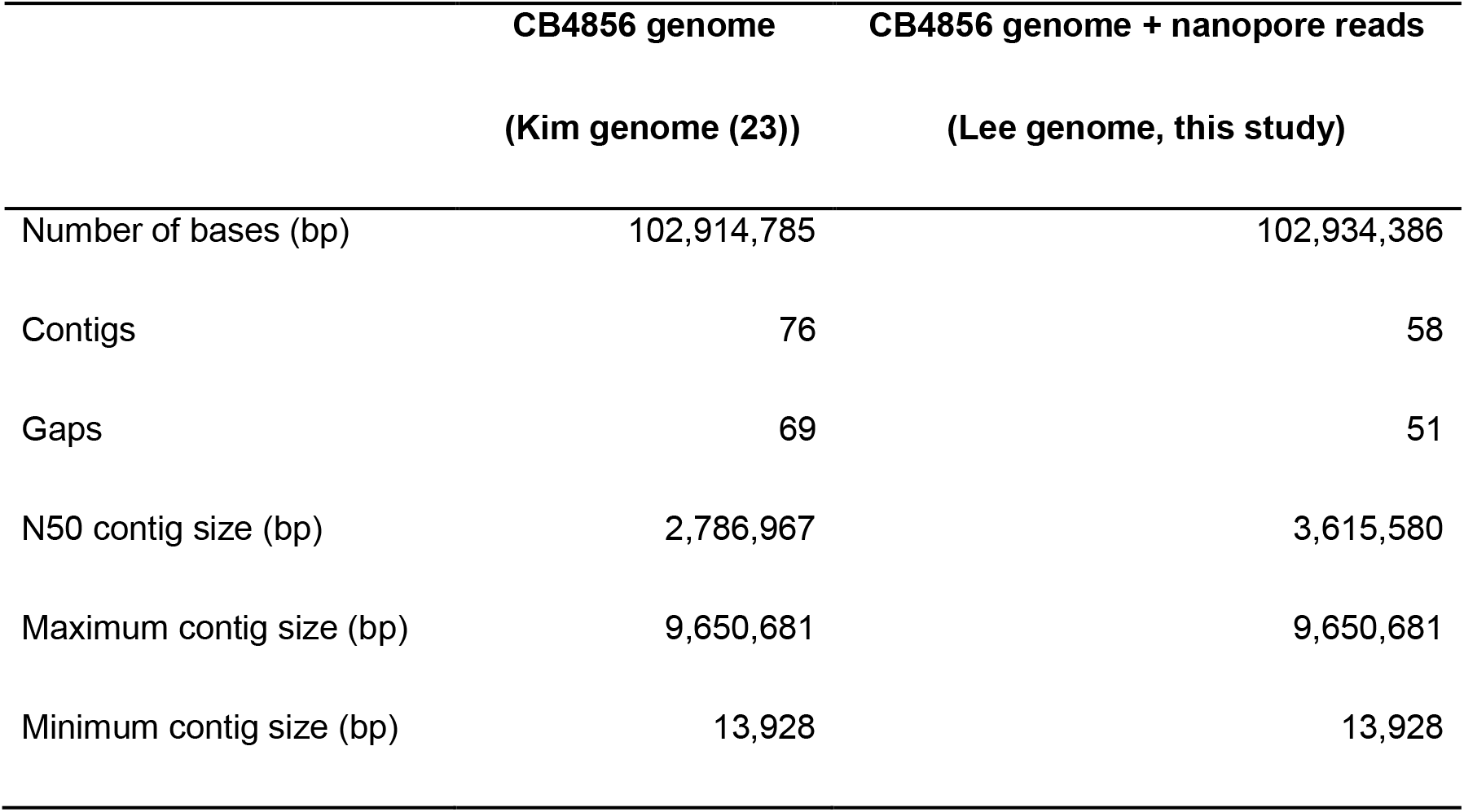
Stats of the Kim genome and the Lee genome for the CB4856 strain

|  | **CB4856 genome** | **CB4856 genome + nanopore reads** |
| --- | --- | --- |
|  | **(Kim genome (23))** | **(Lee genome, this study)** |
| Number of bases (bp) | 102,914,785 | 102,934,386 |
| Contigs | 76 | 58 |
| Gaps | 69 | 51 |
| N50 contig size (bp) | 2,786,967 | 3,615,580 |
| Maximum contig size (bp) | 9,650,681 | 9,650,681 |
| Minimum contig size (bp) | 13,928 | 13,928 |

### Discovery of novel genes in PD1074

After filling nucleotide gaps, we processed long-read RNA sequencing data obtained by a PacBio Iso-Seq platform to identify novel genes using full-length transcripts. First, we generated 14 million raw reads of PD1074 (total 30 Gb, N50 2.7 kb) and processed these reads to obtain 8,630 high-quality, full-length and unique transcripts characterised by non-degraded 5’ end, poly(A) tail and polished sequences (**Figure 1A**, **Supplementary Table S1** and **Supplementary Figure S1C**). These high-quality transcripts corresponded to 6,218 genes (**Figure 1A**) out of which 4,045 contained only known isoforms, but 1,916 genes contained novel isoforms of previously known genes (**Supplementary Table S3**). Total 3,021 novel isoforms were detected, while 177 genes did not match any known gene annotation and were therefore categorised as novel gene candidates (**Supplementary Table S3**). These several thousands of novel isoforms support the hypothesis that our long-read RNA sequencing data were suitable for detecting novel transcripts.

**Figure 1.**
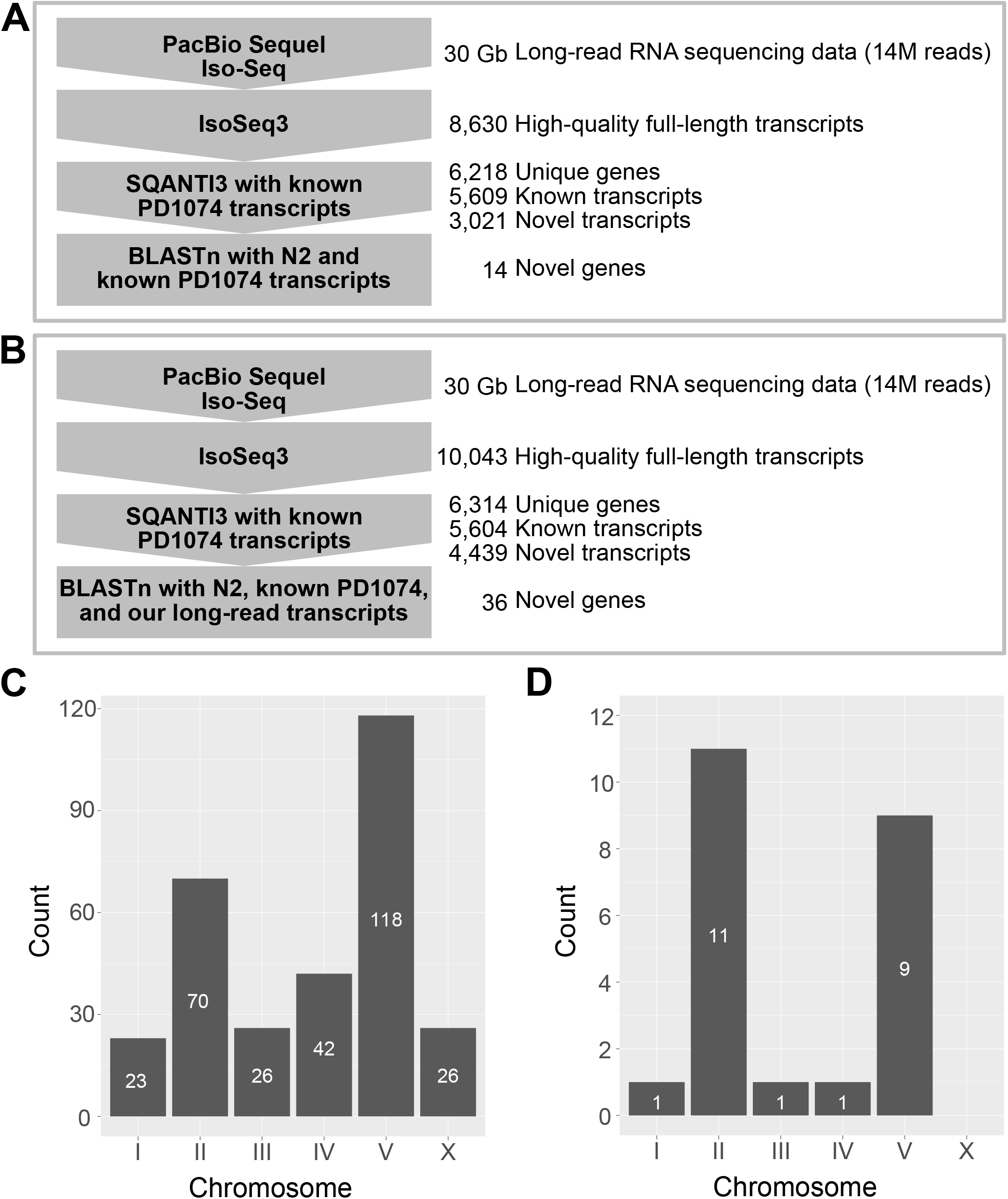
Schematic overview of data analysis to identify novel genes and chromosomal distribution of strain-specific genes. (**A** and **B**) Computational workflows for finding novel genes of (A) PD1074 and (B) CB4856 using long-read RNA sequencing. PacBio Iso-Seq data were processed using IsoSeq3 to produce high-quality full-length transcripts, SQANTI3 to extract novel gene candidates by comparing the transcripts with known PD1074 transcripts and BLASTn to verify the candidates by searching for them in either (A) the N2 and PD1074 known gene databases or (B) the N2 and PD1074 known gene databases supplemented with our long-read PD1074 transcripts database. (**C** and **D**) Chromosomal distribution of (C) PD1074- and (D) CB4856-specific genes.

Among the novel transcripts, 42 were found to correspond to genes belonging to the ribosomal RNA cluster located on the right end of chromosome I (**Supplementary Figure S3**). Their full-length transcripts were longer than those of the previously annotated genes (40), suggesting that our long-read data properly annotated true gene sequences of the previously fragmented genes.

We further validated the 177 novel gene candidates consisting of 182 full-length transcripts to narrow down the list of true novel genes which do not contain any known sequence. We searched for them across the known transcript sequences of PD1074 and its ancestral strain, N2, using BLASTn (**Figure 1A**). We found that 12 non-coding and 2 coding genes did not match any sequence in BLASTn search (**Figure 1A** and **Supplementary Table S4**).

To characterise protein identities of the two coding genes, we searched for their protein sequences in the NCBI protein database and found that a protein sequence encoded by the PB.PD.4262 gene was similar to hypothetical proteins of other *Caenorhabditis* species, but protein sequences encoded by the PB.PD.6031 gene had no significant similarity with any protein sequence within the database (**Supplementary Table S4**). This observation implies that PB.PD.6031 may have been newly born in *C. elegans*.

### Transferring of gene annotation from PD1074 to CB4856

We compared genes between PD1074 and CB4856 by transferring well-annotated gene information from PD1074 to the CB4856 genome (28). Out of 19,954 protein-coding and 26,290 non-coding PD1074 genes, 18,071 (90.6%) protein-coding and 24,992 (95.1%) non-coding genes were transferred to the CB4856 genome (**Supplementary Table S5**). Only 0.2% of the transferred genes were found on different chromosomes and 68.5% of these transcripts were transferred from the PD1074 chromosome V to the CB4856 chromosome II (**Supplementary Table S6**), which is consistent with the previously reported data on this translocated region (23). In total, 6.9% of the PD1074 genes were not transferred and 33% of these transcripts were located on the PD1074 chromosome V (**Supplementary Table S5 and S6**), possibly resulting from small rearrangements in chromosomes V between PD1074 and CB4856 (23,24).

### Discovery of novel genes in CB4856

We additionally generated long-read RNA sequencing data for CB4856 and reported novel transcript information. We processed 14 million raw reads (total 30 Gb, N50 2.6 kb) and obtained 10,043 high-quality, full-length and unique transcripts using PacBio Iso-Seq methods (**Figure 1B**, **Supplementary Table S1** and **Supplementary Figure S1D**). These processed transcripts belonged to 6,314 genes (**Figure 1B** and **Supplementary Table S3**) and, among them, 3,296 genes contained only known isoforms, 2,369 genes contained novel isoforms and 471 genes were categorised as novel gene candidates (**Supplementary Table S3**).

These CB4856 novel gene candidates were further validated using BLAST searches against N2 and PD1074 transcripts, as some of them might exist in the PD1074 genome, but were not transferred. Among the 471 candidates, we determined that 27 protein-coding and 9 non-coding genes, consisting of 54 transcripts, were truly novel as they had no significant matches to any known transcript (**Supplementary Table S7**). The majority of these novel protein-coding genes were similar to other proteins of either *C. elegans* or different *Caenorhabditis* species, but PB.CB.1096, PB.CB.2931 and PB.CB.3084 were found to encode novel proteins, since their protein sequences had no significant similarity with any protein sequence in the NCBI database (**Supplementary Table S7**).

### PAVs between PD1074 and CB4856

Based on these updated transcript data, we identified PAVs differing between PD1074 and CB4856 by comparing PD1074 transcripts to the CB4856 genome to obtain PD1074-specific genes and vice versa for discovering CB4856-specific genes. Considering PD1074, we found that 117 protein-coding and 188 non-coding genes, containing 331 transcripts, were PD1074-specific genes and out of them 239 PAVs were not found in the previous report that used a hybridisation method (**Supplementary Table S8 and S10**) (21). Additionally, 96 genes were found located on the right arm of chromosome V, further supporting the fact that this location can be susceptible to rapid changes both between and within species (**Figure 1C** and **Supplementary Table S8**) (23,24,41). For CB4856, we found 21 protein-coding and two non-coding CB4856-specific genes (**Supplementary Table S9** and **S10**), most of them being localised on chromosomes II and V (**Figure 1D** and **Supplementary Table S9**).

We analysed potential functions of these PAV genes by comparison with known RNAi phenotypes (31). Since *C. elegans* RNAi phenotypes have been studied in the N2 background but not in the CB4856 background, we had an opportunity to investigate only PD1074-specific genes. Among the 117 PD1074-specific protein-coding genes, 11 were reported to have various RNAi phenotypes such as embryonic lethality, reduced brood size, growth variants, oocyte development defects, accumulated cell corpses, protein aggregation variants and hypersensitivity to cadmium and *Bacillus thuringiensis* toxins (**Supplementary Table S8**) (32–38). Although cadmium hypersensitivity was examined in CB4856 as well, this strain showed a similar hypersensitivity response to that of N2 (42).

### Validation of PAVs in other wild strains of *C. elegans* and results of ortholog searches in other *Caenorhabditis* species

In the succeeding experiments, we investigated whether the discovered PAVs were prevalent in natural populations by searching for our PAV sequences in other high-quality genome assemblies among the reported 14 *C. elegans* wild strains (24). We confirmed that 64.3% of our PAVs exhibited presence–absence patterns in other wild strains (**Figure 2A-B** and **Supplementary Table S11** and **S12**). Specifically, all of the CB4856-specific genes derived solely from full-length transcript data had almost identical sequences (more than 90% of identity) in one or more wild strains (4.9 strains on average) (**Figure 2B** and **Supplementary Table S12**). These results suggest the possibility that the discovered CB4856-specific genes were not artefacts that emerged specifically from our CB4856 genome assembly but the genuine genetic variants shared among natural *C. elegans* populations. Conversely, among the total 305 PD1074-specific genes, 208 were found to have almost identical sequences in one or more wild strains (8.4 strains on average), but the remaining 97 genes did not match to any wild strain (**Figure 2A** and **Supplementary Table S11**). It needs to be further verified whether these 97 genes were born during lab-adaptation of PD1074 or they are present in other wild strains not used in our study. Interestingly, 58 PD1074-specific genes and 9 CB4856-specific genes exhibited 10% or more genomic difference in one or more wild strains, as compared to our PAV sequences, suggesting that these genomic regions are still rapidly changing (**Figure 2A-B** and **Supplementary Table S11** and **S12**).

**Figure 2.**
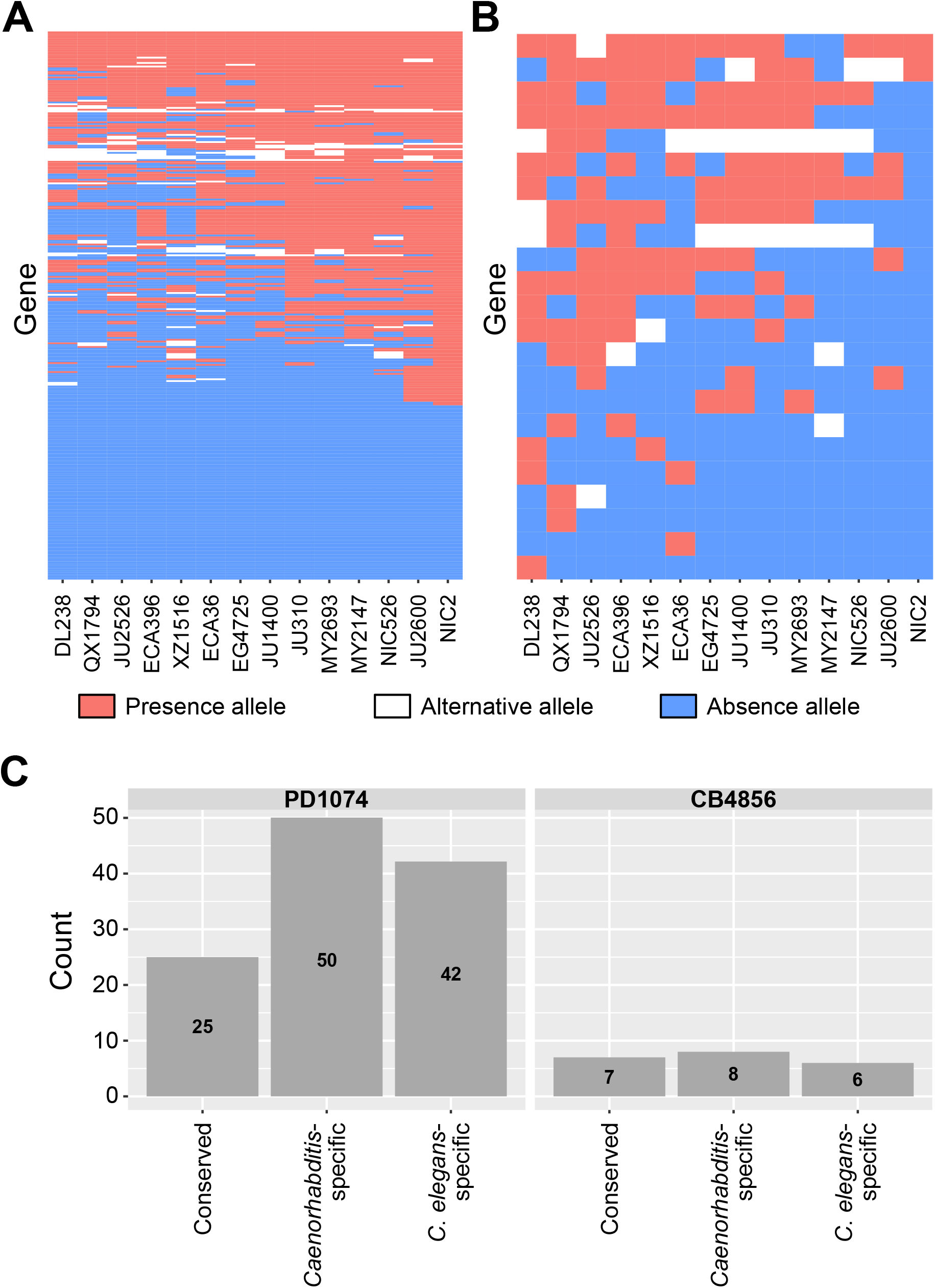
Homology searches for PAV genes to identify alternative alleles and *C. elegans*-specific protein-coding genes. (**A** and **B**) Presence–absence patterns of (A) PD1074-specific and (B) CB4856-specific genes in 14 wild strains. The y-axis represents (A) 306 PD1074-specific and (B) 23 CB4856-specific genes and the x-axis represents wild strains. Red or blue fields indicate if a gene is present or absent in the corresponding strain, respectively. White fields indicate that an allele of a wild strain has substantially different sequences from alleles present in either PD1074 (A) or CB4856 (B) but still possesses > 10% homology sequences. (**C**) Protein homology search results for PAV genes. Each bar means that the discovered genes have similarity with proteins belonging to *C. elegans*, other *Caenorhabditis* species or other organisms.

Thereafter, we examined whether PD1074- and CB4856-specific genes were conserved in other species at the protein level by comparison with the NCBI protein database. Among our PAVs, protein sequences of 93 PD1074- and 15 CB4856-specific genes exhibited at least partial similarity to proteins of other *Caenorhabditis* species and, among these genes, 25 PD1074- and 7 CB4856-specific genes matched to the corresponding protein sequences of other nematodes (**Figure 2C** and **Supplementary Table S8** and **S9**). Intriguingly, proteins encoded by the remaining 42 PD1074- and 6 CB4856-specific genes had no significant similarity with any protein sequence in the database, suggesting that they may have been newly born genes in *C. elegans* (**Figure 2C** and **Supplementary Table S8** and **S9**).

### Alternative alleles of PAV genes may provide evidence of a new gene formation

We assumed that the discovered *C. elegans*-specific PAV genes were recently born through *de novo* gene birth and, if this was correct, other alleles in the *C. elegans* wild strains would have signatures of the rapid gene evolution, since alleles without a specific gene variant (‘absence’) should have gained coding capacities to become ‘presence’ alleles. We analysed whether these 48 PAV genes have alternative alleles, besides presence and absence alleles, among previously published high-quality genome assemblies of 14 *C. elegans* wild strains. We found that 35 PAV genes exhibited only either presence or absence alleles in all 14 genomes, but the remaining 13 PAV genes had alternative alleles that were partially aligned to the corresponding presence alleles (**Figure 3A-C**, **Supplementary Table S13** and **Supplementary Figure S4**).

**Figure 3.**
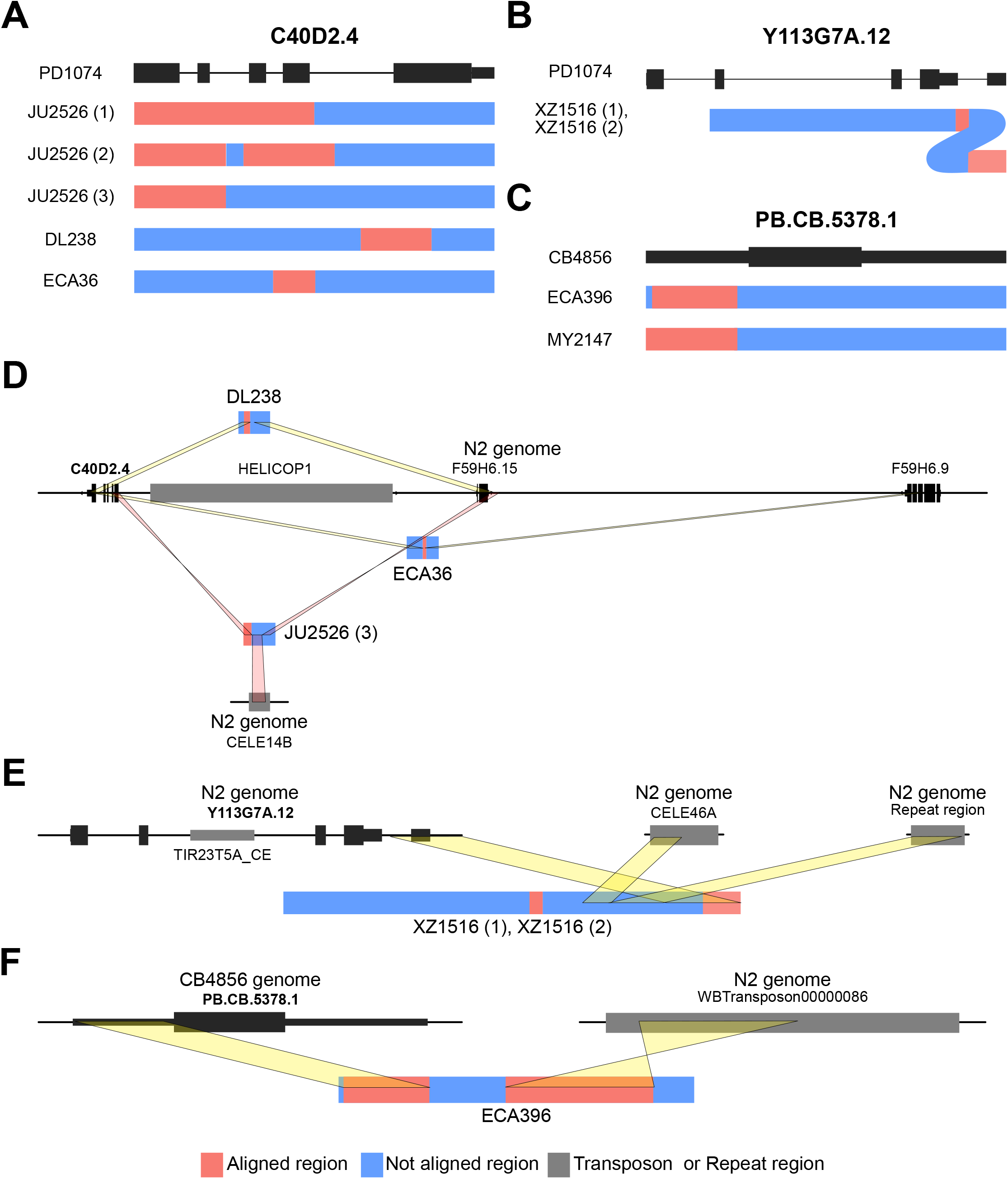
Transposon-mediated allele formation signatures of alternative alleles in two PAV genes. (**A**, **B** and **C**) Schematic allele structure presentation of alternative alleles of (A) C40D2.4, (B) Y113G7A.12 and (C) PB.CB.5378.1. (**D**) Jumping out of the transposon HELICOP1 may produce different alleles of C40D2.4. HELICOP1 transposon (grey block) is located next to the presence allele (left black vertical bars in the central black line) but far from other alternative alleles in DL238 and ECA36. The alternative alleles have partial sequences of C40D2.4 and F59H6.15 or F59H6.9. One of the three alleles of JU2526 contains inverted partial sequences of C40D2.4 and inverted flanking sequences of F59H6.15. These sequences, belonging to two different regions, are connected by another transposon, CELE14B, instead of HELICOP1. (**E**) In the two alternative alleles of Y113G7A.12, 3’ UTR sequences flank the sequences of the CELE46A transposon and a repeat region. (**F**) An alternative allele of PB.CB.5378.1 in ECA396 contains partial original gene sequences and Tc3 family transposon, which do not reside near the original gene in CB4856.

As these alternative alleles may possibly reflect distinguishable events of gene birth or pseudogenisation, we analysed the corresponding presence and alternative alleles in detail. First, the alternative alleles of six genes showed high similarity with other annotated genes (> 31.5% coverage, > 80% identity) and these annotated genes also showed high similarity with the corresponding presence alleles (> 46.7% coverage, > 86% identity), implying that these genes may have been generated through gene duplication and divergence. Among the other seven genes, alternative alleles of F16G10.5, Y46C8AL.11, Y43F8B.22 and PB.CB.5376 were characterised by several small fragments (around 50 bp). These small fragments exhibited no similarity with presence alleles but possessed high similarity to some other sequences in the N2 genome (**Supplementary Figure S4** and **Supplementary Table S13**). We could not find any trace of either duplication or exon shuffling for these alternative alleles, which suggests that their corresponding genes had evolved through extensive indels by unknown mechanisms.

Alternative alleles of the remaining three genes, C40D2.4, Y113G7A.12 and PB.CB.5378.1, exhibited characteristic signatures of the action of transposons. C40D2.4 had five different alternative alleles and its presence allele was found located next to the sequences of the HELICOP1 transposon. No sequence of this transposon flanked the alternative alleles and the sequences of three of them were composed of partial sequences of C40D2.4 and its closely located genes (F59H6.15 or F59H6.9) or non-genic regions possibly derived from transposon jumping out (**Figure 3C**). Moreover, an alternative allele of JU2526 was found to contain partial sequences of another transposon, CELE14B, between inverted partial sequences of C40D2.4 and inverted flanking sequences of F59H6.15 (**Figure 3C**). The presence allele of Y113G7A.12 was found to contain transposon sequences included in the TIR23T5A_CE family. In the two alternative alleles of Y113G7A.12, a transposon sequence was missing and only a partial 3’ UTR region of Y113G7A.12 remained. These partial sequences were also fragmented into two parts and partial sequences of CELE46A transposon and repeat sequences of the N2 genome were found located between the two parts of the 3’ UTR region, suggesting possible evidence of active transposons. PB.CB.5378.1 also had two different alternative alleles. Its presence allele had no transposon sequences, but one alternative allele contained 1.2-kb transposon sequences of the Tc3 family and the other alternative allele contained 80-bp transposon sequences of the CELE46B family (**Figure 3D**). These results suggest the possibility that active transposons may have affected either birth or pseudogenisation of these three genes.

## DISCUSSION

How genes are born and die is an important question in evolutionary genetics, but the answer remains elusive, since this process takes too long to be extensively studied and fully elucidated. Here, we tried to have a glimpse of how genes have evolved by using plenty of genetic resources of *C. elegans* wild strains and long-read sequencing technologies to finely resolve their variations. Specifically, we utilised 48 species-specific PAV genes and alternative alleles of some PAV genes to identify rapid gene evolution snapshots of gene birth and death, since these genes might have been newly born in *C. elegans* and not fixed to the presence alleles. Thirty-five out of these genes did not show any homology at either protein or domain level with other species; therefore, they may not be the genes formed by gene duplication, segmental duplication or whole-genome duplication events, in addition to insertion events of the active transposon (43–45). Unfortunately, we could not find any evidence to understand how these 35 genes have evolved. However, we found that the remaining 13 *C. elegans*-specific protein-coding PAV genes have alternative alleles in other wild isolates. Six out of these genes had similarities to other annotated genes, suggesting that these genes have been generated by gene duplications and changed by mutations. The reason why gene duplication is a major mechanism to produce *C. elegans*-specific genes should be further addressed, as it possibly might happen by chance or from the fact that segmental duplication can produce many genes in a single replication.

On the contrary, alternative alleles of the other seven genes did not exhibit gene duplication signatures, suggesting that they might represent early forms generated through *de novo* gene birth. Three genes, C40D2.4, Y113G7A.12 and PB.CB.5378.1, exhibited either transposon-mediated *de novo* gene birth or pseudogenisation, similarly to new genes that have gained transcription factor functions through transposon insertion, previously reported in a mammalian study (43). However, while this study has shown that active transposons have to be inserted into the existing genes to become DNA-binding motifs, our study shows that the potential for gene birth or death can be achieved through active transposable elements (**Figure 4A-B**). Transposon insertion into a genomic region may add new sequences to the existing non-genic sequences, generating a new gene (**Figure 4A**). In another case, transposon jumping out from a genomic region can lead to a fusion between its surrounding sequences, creating a new gene (**Figure 4B**). The other four genes, which may not result from active transposons, exhibited another gene evolution process that had accumulated small indels. Although the exact sequential process still remains elusive, alternative alleles containing small indels may reveal a *de novo* gene birth process from the non-genic, absence alleles into the genic, presence alleles (or a pseudogenization process, vice versa) (**Figure 4C**).

**Figure 4.**
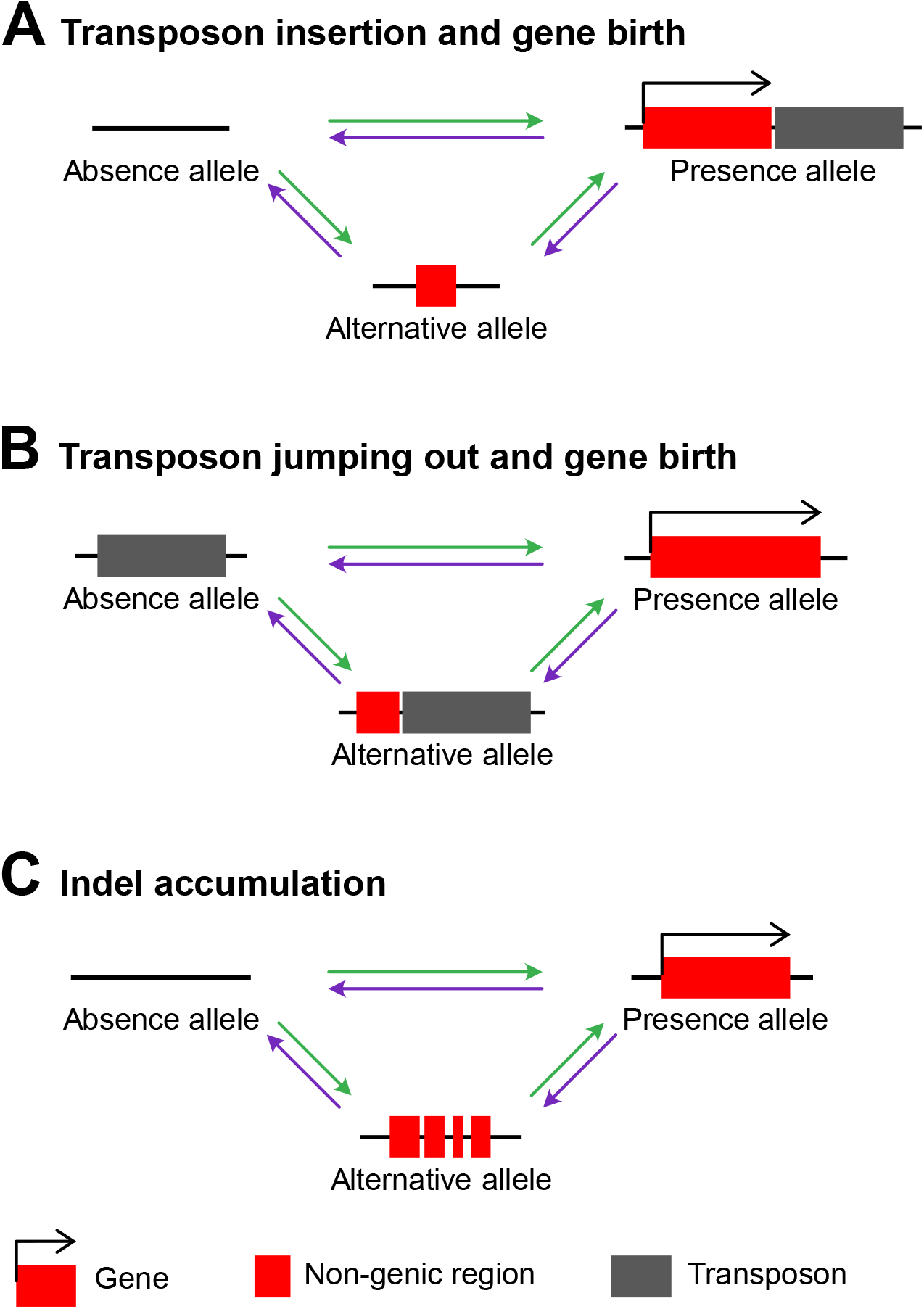
Models of PAV gene evolution. (**A**) A new gene formation or gene loss process by transposon insertion. (**B**) A new gene formation or gene loss process by transposon jumping out. (**C**) A new gene formation or pseudogenisation by small indels. Changes in the nucleotide sequences caused by small indels lead to birth or death of a gene. Red boxes represent genic or pseudogenised genic regions. Black right-angle arrows indicate coding potential of the region. Grey boxes show transposons. Horizontal black lines represent genomic regions. Green arrows represent gene birth processes and purple arrows represent gene loss processes.

Transposons belonging to the TIR23T5A_CE and the TC3 families found in the presence allele of Y113G7A.12 and an alternative allele of PB.CB.5378.1 are DNA transposons known to have ability to be transferred to a new part of a genome through the cut-and-paste mechanism (40,46). HELICOP1 transposon, located next to C40D2.4 in PD1074, is a helitron, which replicates elsewhere in the genome through the rolling circle mechanism (47). Helitrons have also been reported to move to new locations through the cut-and-paste mechanism in maize (48). These cut-and-paste processes cause double-strand breaks in areas from where the transposon has been pulled out and must be repaired by either homologous recombination or non-homologous end-joining (49). In particular, non-homologous end-joining may cause small indels, increase mutation rate of surrounding sequences and/or insert substantially large sequences (50–52). This process may probably contribute to the generation of genes and alternative alleles that we found.

Among PD1074-specific PAV genes, 38.7% were found mainly concentrated on chromosome V, demonstrating that this chromosome is a hotspot for rapid evolution. Our results are highly consistent with those reported previously, implying that chromosomes II and V of N2 contained most of the genes deleted in 12 wild strains (21) and that chromosomes II and V of *C. briggsae* contained lower number of orthologous genes between *C. elegans* and *C. briggsae* genomes (41). Moreover, these characteristics are enriched in the chromosome arm, which is consistent with the previous findings that many core genes are located in the chromosome centre and that new genes are mainly located on the arm (41,53). This suggests that higher density of transposable elements, higher crossover rates or possibly intensive rearrangements in the chromosome arms may have contributed to gene evolution (23,41,54).

Some of the discovered PAV genes could be important for adaptation to different environmental conditions, as they are known to exhibit RNAi phenotypes for important traits such as embryonic lethality and reduced brood size, but their absence alleles still exist in different genetic backgrounds (32–37). Although these results suggest that PAV-related phenotypes can be detected in the CB4856 strain, we could not verify their phenotypes owing to the absence of a genome-wide RNAi or a mutant study on the CB4856 genetic background. One exception was the cadmium-hypersensitive phenotype. However, a phenotypic difference in cadmium hypersensitivity was not observed between N2 and CB4856 (42), implying that the PAV gene may have unknown epistatic or complement genetic relationships. If we take the advantage of powerful reverse genetics tools developed for *C. elegans*, we would be able to define whether PAV-related phenotypic variations exist, and if so, we might be able to find the answer on how PAVs affect such phenotypic variations in different genetic architectures.

Interestingly, we identified new various isoforms using long-read RNA sequencing, despite the fact that gene annotation of *C. elegans* has been updated for over several decades. Among the reported PD1074 long-read transcripts, 35% unique transcripts were categorised as novel isoforms, suggesting that much more isoforms might be present in *C. elegans*, which we may have not been able to detect with conventional approaches. In humans, it is predicted that genes have thousands of isoforms on average (55); therefore, if we succeed to obtain much more full-length transcripts in a simple organism such as *C. elegans*, we would be probably able to identify almost all transcripts, which would allow for further understanding and defining what a gene may represent at the whole species level. Although it is still difficult to grasp whether alternatively spliced isoforms are functional, previous studies showed that the expression of different isoforms may depend on environmental changes, which are known to affect survival and evolution of various organisms (56,57). Thus, it is expected that we can get a deeper insight into this simple organism by elucidating under which condition various isoforms are expressed and what function they have.

In summary, in this study we presented possible processes that may exist between gene birth and death by examining presence, absence and alternative alleles of *C. elegans*-specific genes. Gene duplication and divergence process was found to be a key mechanism for the formation of new genes, but alternative alleles that accumulate small indels were found in some genes, while the other genes have evolved via active transposons. Although the results are still insufficient to fully understand the *de novo* gene birth process, future development of high-quality genomes and genetic models of more diverse wild strains and close relative species of *C. elegans* will enable a precise interpretation of both gene birth and pseudogenisation processes with much higher resolution. It will deepen our understanding of how different organisms acquire and discard genes and how they diverge into different species.

## DATA AVAILABILITY

Our genome assemblies and raw PacBio reads were submitted to the NCBI BioProject database (https://www.ncbi.nlm.nih.gov/bioproject) under accession number PRJNA764925.

## ACKNOWLEDGEMENTS

Author contributions: Conceptualisation, J.K. and B.Y.L.; Methodology, B.Y.L. and J.K.; Formal Analysis, B.Y.L.; Investigation, B.Y.L.; Writing-Original Draft, J.K. and B.Y.L.; Writing-Review & Editing, J.K., B.Y.L., and J.L.; Funding Acquisition, J.L.; Supervision, J.L. and J.K.. Thanks to Chuna Kim for providing the initial idea of this study. Caenorhabditis Genetics Center provided the PD1074 and CB4856 strains.

## SUPPLEMENTARY INFORMATION

**Supplementary Figure S1**. Length distribution of (A) ONT DNA sequencing reads of the CB4856 strain, (B) ONT DNA sequencing reads longer than 20 kb, (C) PacBio Iso-Seq reads of the PD1074 strain and (D) PacBio Iso-Seq reads of the CB4856 strain.

**Supplementary Figure S2**. Dot plots and schematic alignment representations between ONT reads and gap-filled contigs. Each read spans through two contigs at both sides of a corresponding gap.

**Supplementary Figure S3**. Novel transcripts in the rRNA cluster. Forty-two novel transcripts were discovered in the rRNA cluster on chromosome I.

**Supplementary Figure S4**. The alignment pattern of alternative alleles. *C*. *elegans* protein-coding-specific genes exist in wild strains as alternative forms. Red or blue colours indicate aligned or unaligned regions to the corresponding presence allele, respectively.

**Supplementary Table S1**. Raw read stats for long-read DNA and RNA sequencing data.

**Supplementary Table S2**. ONT long-read information used to fill gaps in the previous CB4856 genome.

**Supplementary Table S3**. The stat of genes of PD1074 and CB4856 found by long-read RNA sequencing

**Supplementary Table S4**. List of PD1074 novel genes

**Supplementary Table S5**. LiftOver statistics of gene annotation from PD1074 to CB4856

**Supplementary Table S6**. LiftOver results based on chromosome

**Supplementary Table S7**. List of CB4856 novel genes

**Supplementary Table S8**. Transcript features of PD1074-specific genes

**Supplementary Table S9**. Transcript features of CB4856-specific gene

**Supplementary Table S10**. Stats of PAVs

**Supplementary Table S11**. PD1074-specific gene distributions in 14 wild isolates

**Supplementary Table S12**. CB4856-specific gene distribution in 14 wild isolates

**Supplementary Table S13**. The loci of alternative alleles

## REFERENCES

1. Jacob, F. (1977) Evolution and tinkering. Science, 196, 1161–1166.

2. Dennis, M.Y. and Eichler, E.E. (2016) Human adaptation and evolution by segmental duplication. Curr Opin Genet Dev, 41, 44–52.

3. Marlétaz, F., Firbas, P.N., Maeso, I., Tena, J.J., Bogdanovic, O., Perry, M., Wyatt, C.D.R., de la Calle-Mustienes, E., Bertrand, S., Burguera, D. et al. (2018) Amphioxus functional genomics and the origins of vertebrate gene regulation. Nature, 564, 64–70.

4. Van Oss, S.B. and Carvunis, A.R. (2019) De novo gene birth. Plos Genetics, 15.

5. Begun, D.J., Lindfors, H.A., Kern, A.D. and Jones, C.D. (2007) Evidence for de novo evolution of testis-expressed genes in the Drosophila yakuba/Drosophila erecta clade. Genetics, 176, 1131–1137.

6. Levine, M.T., Jones, C.D., Kern, A.D., Lindfors, H.A. and Begun, D.J. (2006) Novel genes derived from noncoding DNA in Drosophila melanogaster are frequently X-linked and exhibit testis-biased expression. Proc Natl Acad Sci U S A, 103, 9935–9939.

7. Begun, D.J., Lindfors, H.A., Thompson, M.E. and Holloway, A.K. (2006) Recently evolved genes identified from Drosophila yakuba and D. erecta accessory gland expressed sequence tags. Genetics, 172, 1675–1681.

8. Carvunis, A.R., Rolland, T., Wapinski, I., Calderwood, M.A., Yildirim, M.A., Simonis, N., Charloteaux, B., Hidalgo, C.A., Barbette, J., Santhanam, B. et al. (2012) Proto-genes and de novo gene birth. Nature, 487, 370–374.

9. Takahashi-Kariyazono, S., Sakai, K. and Terai, Y. (2020) Presence–absence polymorphisms of single-copy genes in the stony coral Acropora digitifera. BMC Genomics, 21, 158.

10. Winzer, T., Gazda, V., He, Z., Kaminski, F., Kern, M., Larson, T.R., Li, Y., Meade, F., Teodor, R., Vaistij, F.E. et al. (2012) A Papaver somniferum 10-gene cluster for synthesis of the anticancer alkaloid noscapine. Science, 336, 1704–1708.

11. Gabur, I., Chawla, H.S., Lopisso, D.T., von Tiedemann, A., Snowdon, R.J. and Obermeier, C. (2020) Gene presence–absence variation associates with quantitative Verticillium longisporum disease resistance in Brassica napus. Sci Rep, 10, 4131.

12. Jiang, L., Lv, Y., Li, T., Zhao, H. and Zhang, T. (2015) Identification and characterization of presence/absence variation in maize genotype Mo17. Genes Genom, 37, 503–515.

13. Rosa, R.D., Alonso, P., Santini, A., Vergnes, A. and Bachere, E. (2015) High polymorphism in big defensin gene expression reveals presence–absence gene variability (PAV) in the oyster Crassostrea gigas. Dev Comp Immunol, 49, 231–238.

14. Shen, J., Araki, H., Chen, L., Chen, J.Q. and Tian, D. (2006) Unique evolutionary mechanism in R-genes under the presence/absence polymorphism in Arabidopsis thaliana. Genetics, 172, 1243–1250.

15. Calcino, A.D., Kenny, N.J. and Gerdol, M. (2021) Single individual structural variant detection uncovers widespread hemizygosity in molluscs. Philos Trans R Soc Lond B Biol Sci, 376, 20200153.

16. Gao, L., Gonda, I., Sun, H., Ma, Q., Bao, K., Tieman, D.M., Burzynski-Chang, E.A., Fish, T.L., Stromberg, K.A., Sacks, G.L. et al. (2019) The tomato pan-genome uncovers new genes and a rare allele regulating fruit flavor. Nat Genet, 51, 1044–1051.

17. Liu, Y., Du, H., Li, P., Shen, Y., Peng, H., Liu, S., Zhou, G.A., Zhang, H., Liu, Z., Shi, M. et al. (2020) Pan-Genome of Wild and Cultivated Soybeans. Cell, 182, 162–176 e113.

18. Li, C., Xiang, X., Huang, Y., Zhou, Y., An, D., Dong, J., Zhao, C., Liu, H., Li, Y., Wang, Q. et al. (2020) Long-read sequencing reveals genomic structural variations that underlie creation of quality protein maize. Nat Commun, 11, 17.

19. Cook, D.E., Zdraljevic, S., Roberts, J.P. and Andersen, E.C. (2017) CeNDR, the Caenorhabditis elegans natural diversity resource. Nucleic Acids Res, 45, D650–D657.

20. Crombie, T.A., Zdraljevic, S., Cook, D.E., Tanny, R.E., Brady, S.C., Wang, Y., Evans, K.S., Hahnel, S., Lee, D., Rodriguez, B.C. et al. (2019) Deep sampling of Hawaiian Caenorhabditis elegans reveals high genetic diversity and admixture with global populations. Elife, 8.

21. Maydan, J.S., Lorch, A., Edgley, M.L., Flibotte, S. and Moerman, D.G. (2010) Copy number variation in the genomes of twelve natural isolates of Caenorhabditis elegans. BMC Genomics, 11, 62.

22. Yoshimura, J., Ichikawa, K., Shoura, M.J., Artiles, K.L., Gabdank, I., Wahba, L., Smith, C.L., Edgley, M.L., Rougvie, A.E., Fire, A.Z. et al. (2019) Recompleting the Caenorhabditis elegans genome. Genome Res, 29, 1009–1022.

23. Kim, C., Kim, J., Kim, S., Cook, D.E., Evans, K.S., Andersen, E.C. and Lee, J. (2019) Long-read sequencing reveals intra-species tolerance of substantial structural variations and new subtelomere formation in C. elegans. Genome Res, 29, 1023–1035.

24. Lee, D., Zdraljevic, S., Stevens, L., Wang, Y., Tanny, R.E., Crombie, T.A., Cook, D.E., Webster, A.K., Chirakar, R., Baugh, L.R. et al. (2021) Balancing selection maintains hyperdivergent haplotypes in Caenorhabditis elegans. Nat Ecol Evol.

25. Li, H. (2018) Minimap2: pairwise alignment for nucleotide sequences. Bioinformatics, 34, 3094–3100.

26. Li, H., Handsaker, B., Wysoker, A., Fennell, T., Ruan, J., Homer, N., Marth, G., Abecasis, G., Durbin, R. and Genome Project Data Processing, S. (2009) The Sequence Alignment/Map format and SAMtools. Bioinformatics, 25, 2078–2079.

27. Marcais, G., Delcher, A.L., Phillippy, A.M., Coston, R., Salzberg, S.L. and Zimin, A. (2018) MUMmer4: A fast and versatile genome alignment system. PLoS Comput Biol, 14, e1005944.

28. Navarro Gonzalez, J., Zweig, A.S., Speir, M.L., Schmelter, D., Rosenbloom, K.R., Raney, B.J., Powell, C.C., Nassar, L.R., Maulding, N.D., Lee, C.M. et al. (2021) The UCSC Genome Browser database: 2021 update. Nucleic Acids Res, 49, D1046–D1057.

29. Tardaguila, M., de la Fuente, L., Marti, C., Pereira, C., Pardo-Palacios, F.J., Del Risco, H., Ferrell, M., Mellado, M., Macchietto, M., Verheggen, K. et al. (2018) SQANTI: extensive characterization of long-read transcript sequences for quality control in full-length transcriptome identification and quantification. Genome Res.

30. Marchler-Bauer, A., Bo, Y., Han, L., He, J., Lanczycki, C.J., Lu, S., Chitsaz, F., Derbyshire, M.K., Geer, R.C., Gonzales, N.R. et al. (2017) CDD/SPARCLE: functional classification of proteins via subfamily domain architectures. Nucleic Acids Res, 45, D200–D203.

31. Harris, T.W., Arnaboldi, V., Cain, S., Chan, J., Chen, W.J., Cho, J., Davis, P., Gao, S., Grove, C.A., Kishore, R. et al. (2020) WormBase: a modern Model Organism Information Resource. Nucleic Acids Res, 48, D762–D767.

32. Fernandez, A.G., Gunsalus, K.C., Huang, J., Chuang, L.S., Ying, N., Liang, H.L., Tang, C., Schetter, A.J., Zegar, C., Rual, J.F. et al. (2005) New genes with roles in the C. elegans embryo revealed using RNAi of ovary-enriched ORFeome clones. Genome Res, 15, 250–259.

33. Rual, J.F., Ceron, J., Koreth, J., Hao, T., Nicot, A.S., Hirozane-Kishikawa, T., Vandenhaute, J., Orkin, S.H., Hill, D.E., van den Heuvel, S. et al. (2004) Toward improving Caenorhabditis elegans phenome mapping with an ORFeome-based RNAi library. Genome Res, 14, 2162–2168.

34. Sakaki, K., Yoshina, S., Shen, X., Han, J., DeSantis, M.R., Xiong, M., Mitani, S. and Kaufman, R.J. (2012) RNA surveillance is required for endoplasmic reticulum homeostasis. Proc Natl Acad Sci U S A, 109, 8079–8084.

35. Cui, Y., McBride, S.J., Boyd, W.A., Alper, S. and Freedman, J.H. (2007) Toxicogenomic analysis of Caenorhabditis elegans reveals novel genes and pathways involved in the resistance to cadmium toxicity. Genome Biol, 8, R122.

36. Green, R.A., Kao, H.L., Audhya, A., Arur, S., Mayers, J.R., Fridolfsson, H.N., Schulman, M., Schloissnig, S., Niessen, S., Laband, K. et al. (2011) A high-resolution C. elegans essential gene network based on phenotypic profiling of a complex tissue. Cell, 145, 470–482.

37. Zullig, S., Neukomm, L.J., Jovanovic, M., Charette, S.J., Lyssenko, N.N., Halleck, M.S., Reutelingsperger, C.P., Schlegel, R.A. and Hengartner, M.O. (2007) Aminophospholipid translocase TAT-1 promotes phosphatidylserine exposure during C. elegans apoptosis. Curr Biol, 17, 994–999.

38. Kao, C.Y., Los, F.C., Huffman, D.L., Wachi, S., Kloft, N., Husmann, M., Karabrahimi, V., Schwartz, J.L., Bellier, A., Ha, C. et al. (2011) Global functional analyses of cellular responses to pore-forming toxins. PLoS Pathog, 7, e1001314.

39. Kent, W.J. (2002) BLAT--the BLAST-like alignment tool. Genome Res, 12, 656–664.

40. Consortium, C.e.S. (1998) Genome sequence of the nematode C. elegans: a platform for investigating biology. Science, 282, 2012–2018.

41. Stein, L.D., Bao, Z., Blasiar, D., Blumenthal, T., Brent, M.R., Chen, N., Chinwalla, A., Clarke, L., Clee, C., Coghlan, A. et al. (2003) The genome sequence of Caenorhabditis briggsae: a platform for comparative genomics. PLoS Biol, 1, E45.

42. Evans, K.S., Brady, S.C., Bloom, J.S., Tanny, R.E., Cook, D.E., Giuliani, S.E., Hippleheuser, S.W., Zamanian, M. and Andersen, E.C. (2018) Shared Genomic Regions Underlie Natural Variation in Diverse Toxin Responses. Genetics, 210, 1509–1525.

43. Cosby, R.L., Judd, J., Zhang, R., Zhong, A., Garry, N., Pritham, E.J. and Feschotte, C. (2021) Recurrent evolution of vertebrate transcription factors by transposase capture. Science, 371.

44. Crow, K.D. and Wagner, G.P. (2006) What Is the Role of Genome Duplication in the Evolution of Complexity and Diversity? Mol Biol Evol, 23, 887–892.

45. Meyer, A. and Schartl, M. (1999) Gene and genome duplications in vertebrates: the one-to-four (-to-eight in fish) rule and the evolution of novel gene functions. Current Opinion in Cell Biology, 11, 699–704.

46. van Luenen, H.G., Colloms, S.D. and Plasterk, R.H. (1994) The mechanism of transposition of Tc3 in C. elegans. Cell, 79, 293–301.

47. Kapitonov, V.V. and Jurka, J. (2007) Helitrons on a roll: eukaryotic rolling-circle transposons. Trends Genet, 23, 521–529.

48. Li, Y. and Dooner, H.K. (2009) Excision of Helitron transposons in maize. Genetics, 182, 399–402.

49. Krasileva, K.V. (2019) The role of transposable elements and DNA damage repair mechanisms in gene duplications and gene fusions in plant genomes. Curr Opin Plant Biol, 48, 18–25.

50. Wicker, T., Yu, Y., Haberer, G., Mayer, K.F., Marri, P.R., Rounsley, S., Chen, M., Zuccolo, A., Panaud, O., Wing, R.A. et al. (2016) DNA transposon activity is associated with increased mutation rates in genes of rice and other grasses. Nat Commun, 7, 12790.

51. Gorbunova, V. and Levy, A.A. (1997) Non-homologous DNA end joining in plant cells is associated with deletions and filler DNA insertions. Nucleic Acids Res, 25, 4650–4657.

52. Kim, C., Sung, S., Kim, J. and Lee, J. (2020) Repair and Reconstruction of Telomeric and Subtelomeric Regions and Genesis of New Telomeres: Implications for Chromosome Evolution. Bioessays, 42, e1900177.

53. Prabh, N., Roeseler, W., Witte, H., Eberhardt, G., Sommer, R.J. and Rodelsperger, C. (2018) Deep taxon sampling reveals the evolutionary dynamics of novel gene families in Pristionchus nematodes. Genome Res, 28, 1664–1674.

54. Woodruff, G.C. and Teterina, A.A. (2020) Degradation of the Repetitive Genomic Landscape in a Close Relative of Caenorhabditis elegans. Mol Biol Evol, 37, 2549–2567.

55. Sedlazeck, F.J., Lee, H., Darby, C.A. and Schatz, M.C. (2018) Piercing the dark matter: bioinformatics of long-range sequencing and mapping. Nat Rev Genet, 19, 329–346.

56. Trevisan, G.L., Oliveira, E.H., Peres, N.T., Cruz, A.H., Martinez-Rossi, N.M. and Rossi, A. (2011) Transcription of Aspergillus nidulans pacC is modulated by alternative RNA splicing of palB. FEBS Lett, 585, 3442–3445.

57. Nilsen, T.W. and Graveley, B.R. (2010) Expansion of the eukaryotic proteome by alternative splicing. Nature, 463, 457–463.

