## Supplementary figures 1-4 for "Intraspecific *de novo* gene birth revealed by presence–absence variant genes in *Caenorhabditis elegans*"

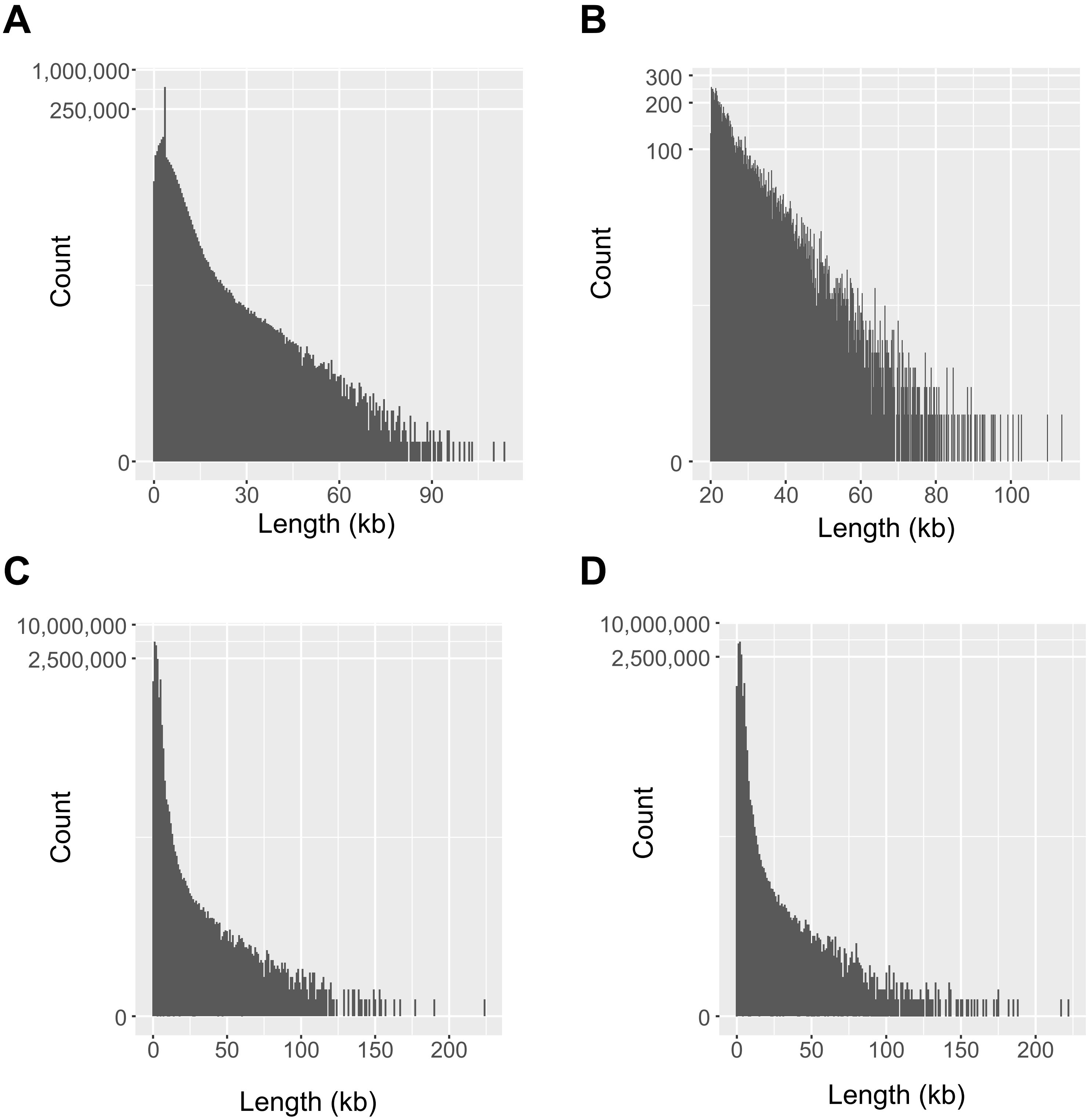


**Supplementary Figure S1**. Length distribution of (**A**) ONT DNA sequencing reads of the CB4856 strain, (**B**) ONT DNA sequencing reads longer than 20 kb, (**C**) PacBio Iso-Seq reads of the PD1074 strain and (**D**) PacBio Iso-Seq reads of the CB4856 strain

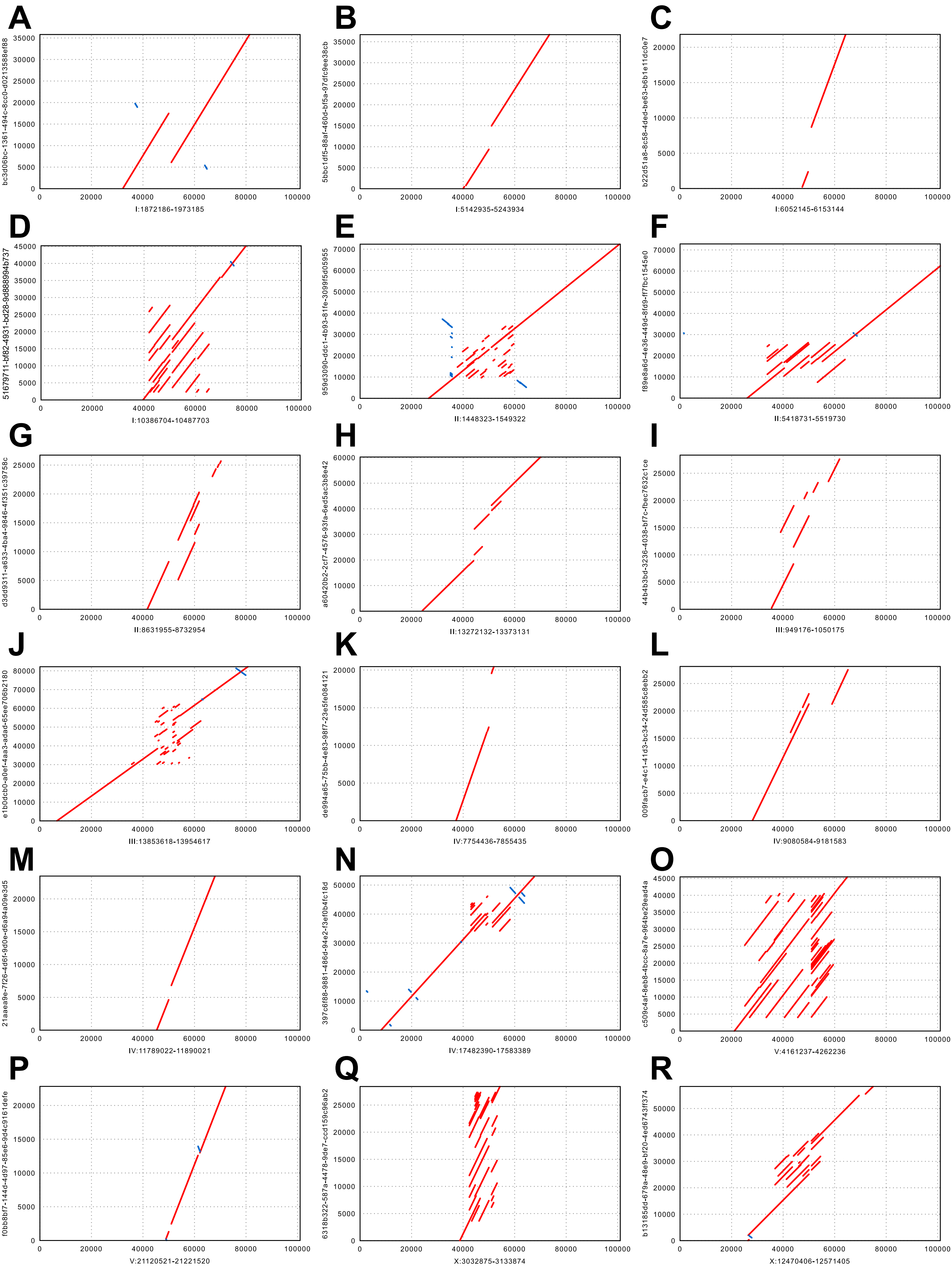


**Supplementary Figure S2**. Dot plots and schematic alignment representations between ONT reads and gap-filled contigs. Each read spans through two contigs at both sides of a corresponding gap.

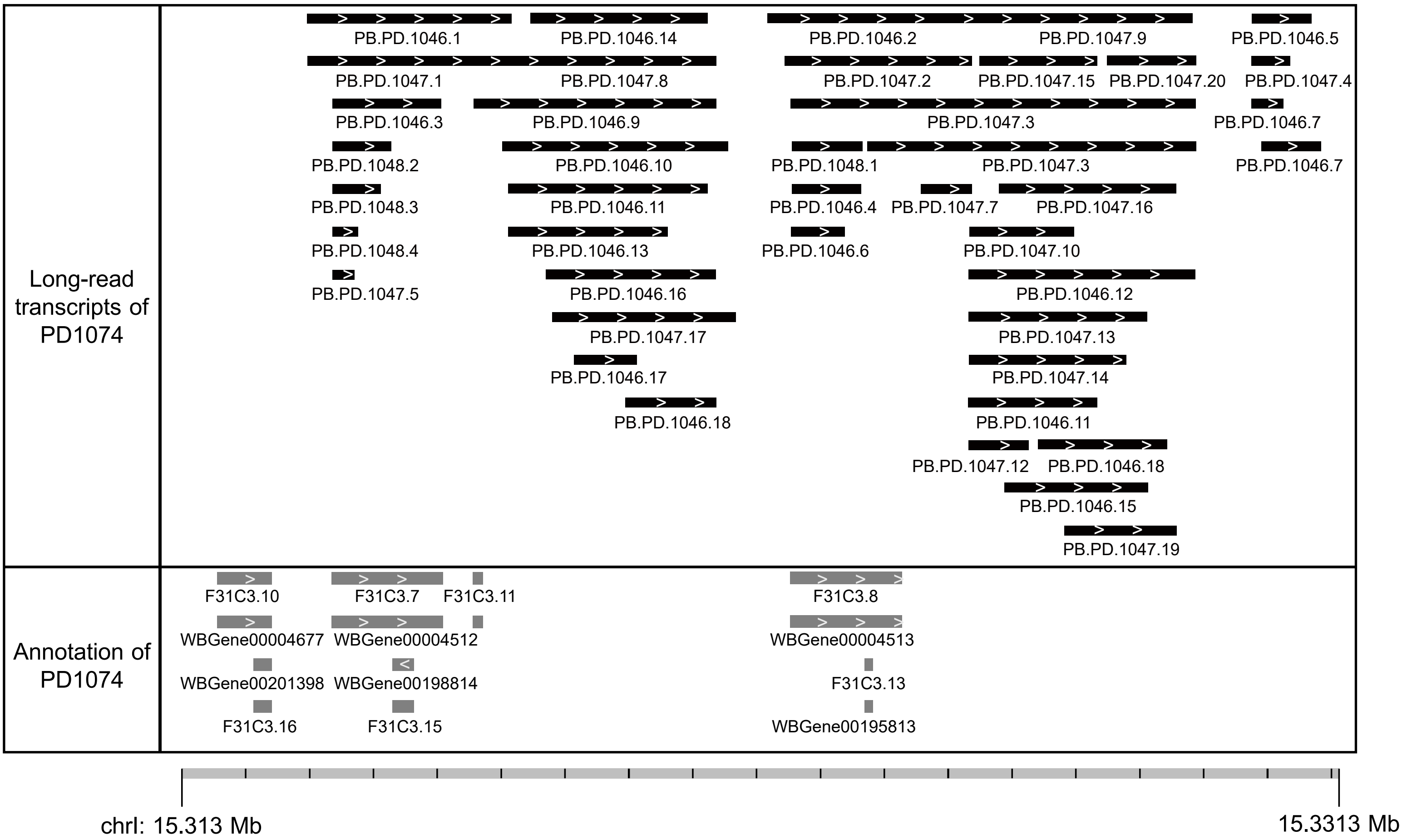


**Supplementary Figure S3**. Novel transcripts in the rRNA cluster. Forty-two novel transcripts were discovered in the rRNA cluster on chromosome I.

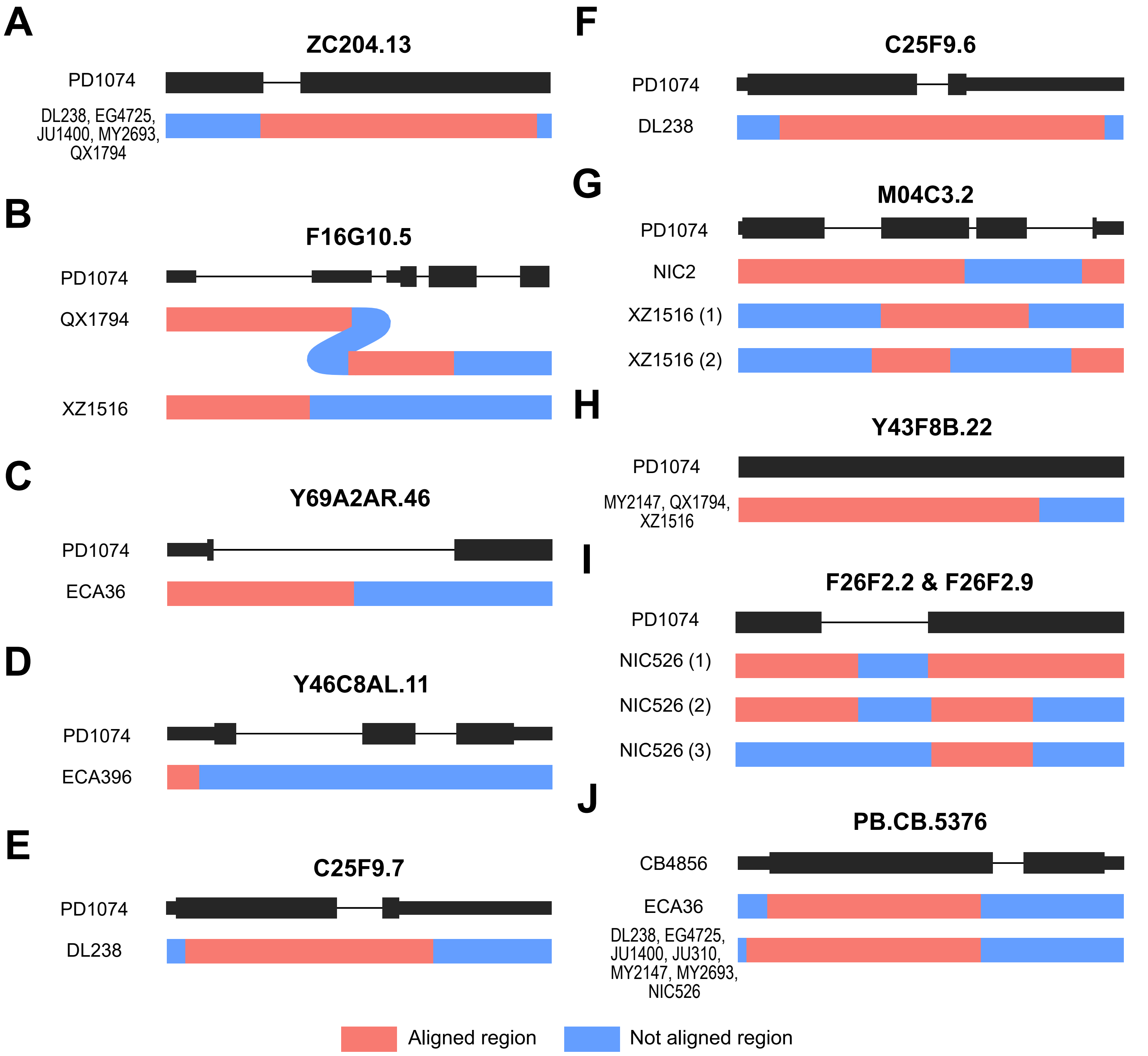


**Supplementary Figure S4**. The alignment pattern of alternative alleles. *C. elegans* protein-coding-specific genes exist in wild strains as alternative forms. Red or blue colours indicate aligned or unaligned regions to the corresponding presence allele, respectively.
